# Animal personality adds complexity to the processes of adaptive divergence and speciation

**DOI:** 10.1101/2020.03.02.971614

**Authors:** Quentin J.B. Horta-Lacueva, David Benhaïm, Michael B. Morrissey, Sigurður S. Snorrason, Kalina H. Kapralova

## Abstract

Divergent selection is a powerful driver of speciation and has been widely studied in relation to the physical characters of organisms. Because evolution of behavioural traits may contribute to evolutionary processes, we explored how consistent variation in behaviours may affect the process of adaptive divergence and speciation. We studied whether two sympatric morphs of Arctic charr (*Salvelinus alpinus*) have recently evolved genetically-based differences in personality that conform to their respective ecological niches, and whether these differences contribute to reproductive isolation by generating maladaptive hybrid behaviours. Studying three aspects of behavioural variation (average trait value, consistent individual differences and trait covariance), we assessed the sociality and risk-taking propensity of hybrid and pure-morph offspring reared in common conditions. Contrary to expectations, the two morphs did not differ in the average values of these traits but showed different behavioural syndromes (trait covariances). While the hybrids did not differ from either morph in their average behavioural responses, they showed less individual consistency in these behaviours and a different set of behavioural syndromes. Differences between morphs and their hybrids in other behavioural aspects than their average behavioural responses suggest that our understanding of speciation processes can benefit from an integrative view of behavioural variation.

## Introduction

A variety of behavioural traits are now widely recognized as playing a key role in the evolution of animal populations through natural selection [1–3]. Traits such as aggressivity, sociality and boldness (the propensity for taking risks) are well studied for their implications on foraging success, predator avoidance, vulnerability to parasites, breeding success, and life-history strategies in a wide range of taxa (e.g. arthropods, vertebrates, molluscs, sea anemones) [4–9]. Major advances in our understanding of the role of behaviour in evolution have been recently achieved through studies of behavioural differences at the individual level, and focusing on individual consistency, that is, animal personality [10–12]. Behavioural differences among populations are now considered as more than adaptive mean values surrounded by the noise of individual variance and can be studied under different aspects, such as (i) the group level, average values of a given behavioural trait, (ii) the consistent differences between individuals in this trait (personality *per se*) and (iii) the covariations between this trait and others (behavioural syndromes) [4]. Because of their genetic bases and their effects on fitness, personality and behavioural syndromes are considered to be important drivers of adaptive divergence and speciation [3,13,14]. Considering classical models of speciation [15,16], divergent selection may indeed result in the segregation of behavioural types between different fitness optima. Reproductive isolation (such as selection against intermediate or transgressive hybrids) could then develop as a by-product, which may in turn lead to further divergence [17]. Despite the recent conceptual advances about the importance of personality in the processes of adaptive divergence and speciation, empirical studies directly investigating it are still lacking [14]. Information is especially lacking regarding the behavioural phenotype of hybrids and their contribution to reproductive isolation [18].

Here we explore whether behaviour as considered under the three aspects described above (average trait value, consistent differences between individuals and trait covariance) can influence the evolutionary processes of divergence and speciation. First, contrasting ecological conditions can generate different fitness optima that favour the differentiation of populations in the average values of a behavioural response (*i.e*. “behavioural adjustment”, Figure 1a) [19]. Second, contrasting environmental variables such as the predictability of a food resource or predation risk can determine the benefit of behavioural consistency over plasticity, thus affecting the level of consistent differences between individuals (i.e. personality *per se*) [20] – defined as behavioural “homogenisation” vs. “diversification” in [19] (Figure 1). Finally, because of genetic constraints [21,22], functional trade-offs [23] and multidimensional selection [24], covariances between traits can be important determinants of fitness and one may therefore expect personality syndromes to be shaped differently between diverging populations [25]. These three aspects of behavioural variation could therefore affect the build-up of reproductive isolation if hybrids present either (i) disadvantageous average values in personality traits, (ii) a loss in behavioural consistency (personality breakdown) or (iii) maladaptive combinations of these traits (syndrome breakdown).

**Figure 1.**
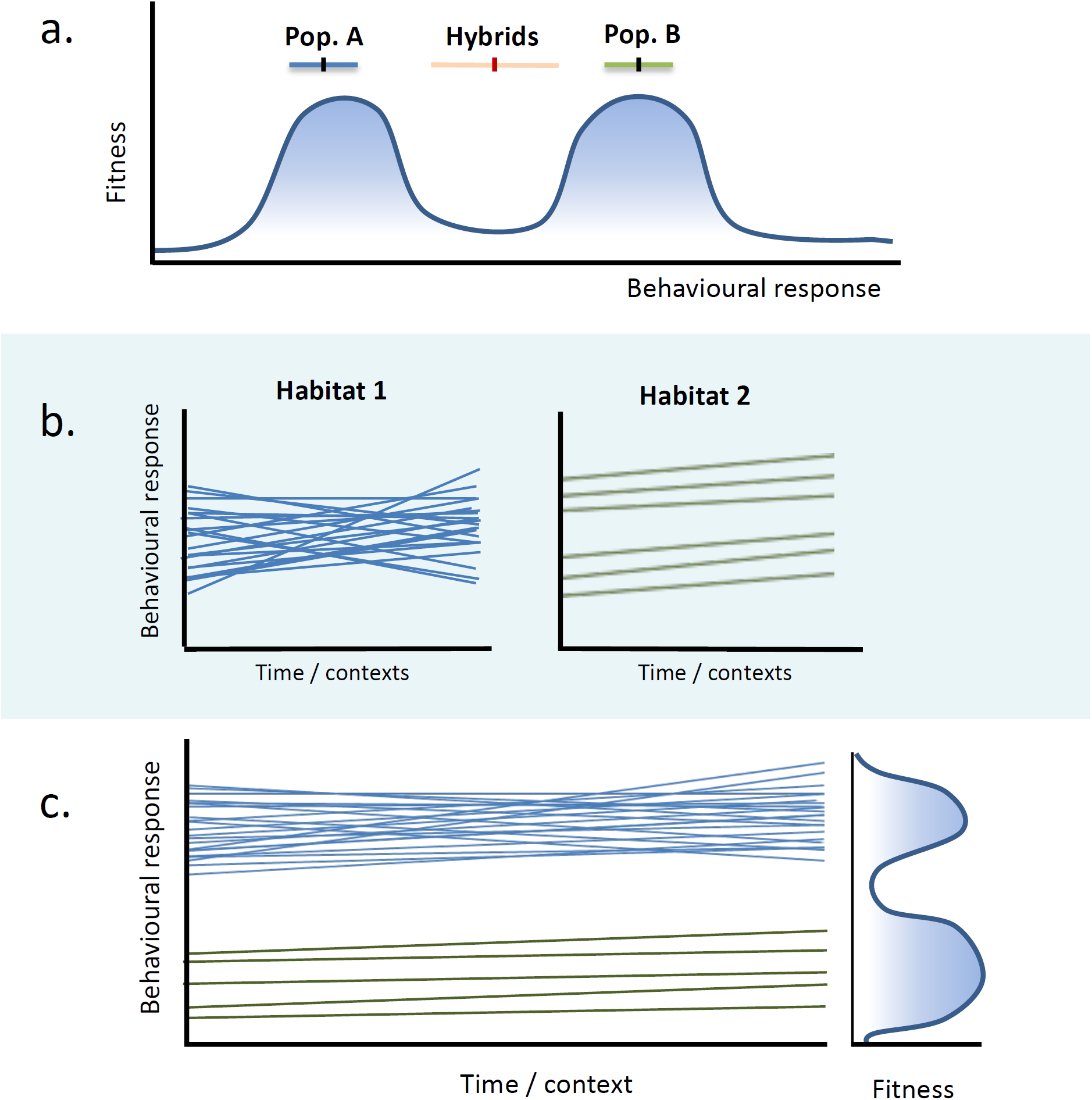
Evolutionary mechanism through which behavioural variation is involved in adaptive divergence. (a) Average behavioural response can diverge between populations as any classic trait under divergent selection; here represented in a rugged-adaptive landscape model [15]. Reproductive isolation can build-up as the intermediate or transgressive behavioural responses of hybrids represent a selective disadvantage. (b) Different selection regimes can also affect the level of consistent behavioural differences between individuals (*i.e.* personality) across environments. This can be visualized under a reaction norm approach. While in this example identical average values are favoured in two environments, specialized behavioural types are favoured in Environment 2. (c) Complex patterns of adaptive divergence may therefore arise when considering the two aspects of behavioural variations described in (a) and (b).

Postglacial lakes hosting different varieties or morphs of freshwater fish are particularly valuable biological systems offering a glimpse of early stages of divergence [26]. These lakes often contain sympatric populations, which facilitate the study of divergent selection by limiting the effects of geographical barriers on gene flow [27]. The evolution of these systems has been described under the framework of resource polymorphism, where a few fish species colonized recently de-glaciated lakes offering a variety of unoccupied ecological niches, thus promoting the emergence of different sympatric morphs [28]. These morphs usually segregate between the benthic and the limnetic habitats and are characterised by various levels of reproductive isolation [29,30]. The Arctic charr from the Icelandic lake Thingvallavatn presents an extreme and rapid case of such divergence that resulted in the emergence of four lake-locked morphs which are evolving (at least in their current state) in sympatry (Figure 2a).

**Figure 2.**
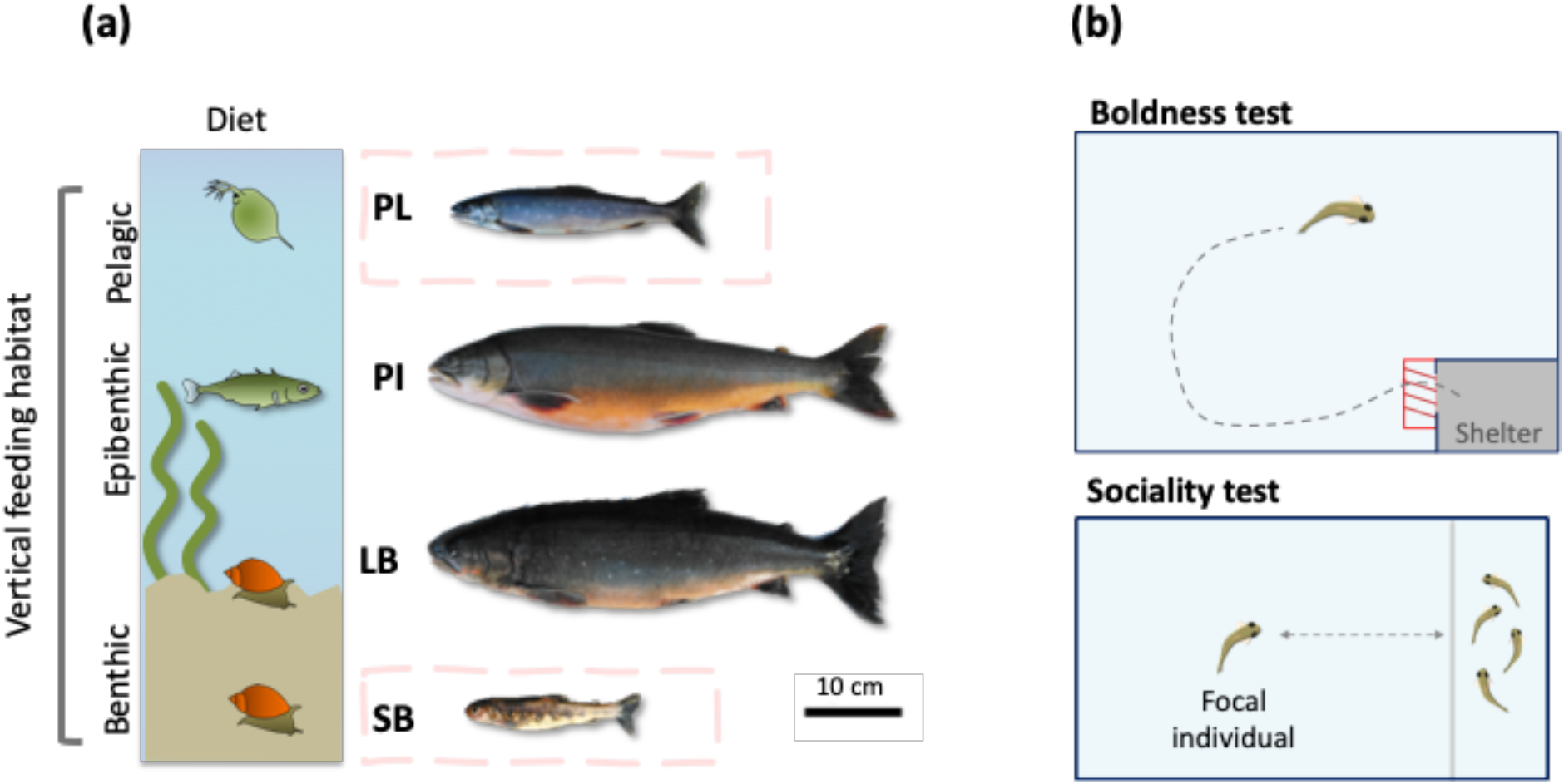
Studied morphs and experimental system. (a) The four morphs of Thingvallavatn with their foraging habitat and main prey depicted on the left side (from top to bottom: planktonic crustacean, fish, freshwater snail). All four specimens were captured together on the same site. PL: Planktivorous, PI: Piscivorous, LB: Large-benthic, SB: Small-benthic charr. The two focus morphs of our study (PL and SB) are higlited with dashed lines. (b) Schematic view of the two experimental arenas used to measure the boldness and sociality traits (aerial view). In the top arena, the dashed zone represents the near entrance of the shelter that was used to describe an additional personality axis defined as the “doormat” trait. Dimensions are shown in Figure S1.

We focus here on two of the four morphs in Thingvallavatn, the “small-benthic” (SB) and the “planktivorous” (PL) charrs. The SB charr live in the stony littoral zone of the lake and forage on benthic invertebrates, mainly the snail *Radix peregra* and chironomid larvae. In this habitat they use their small size to manoeuvre amongst the lava stones in order to access food and seek shelter from predation. The PL charr utilize the pelagic zone of the lake and feed on zooplankton and emerging chironomids. The spawning seasons of SB- and PL-charr overlap (the spawning season of SB charr encompassing the one of PL charr) and these two morphs appear to also share their spawning locations [31]. Estimates of gene flow between the two charr are however very low [32] and individuals of intermediate morphology are rarely observed in spite of the ease to generate mature first-generation hybrids (F_1_) in captivity [31,33]. These observations suggest that selection against hybrids may at least to some extent contribute to the reproductive isolation of the two morphs.

Given the ecological differences between the small benthic and planktivorous morphs, we expected them to have evolved differences in boldness and sociality, two personality traits known to be often related to fitness [6,9]. These differences should be reflected by changes in the three behavioural aspects discussed above (see our predictions in Table 1). Briefly, we expected the PL charr to have evolved personality traits favouring the formation of shoals. Shoaling formation is well understood to be advantageous in open habitats with low physical complexity [34,35]. During the summer and autumn, juvenile and adult PL charr forage in open water environments where they spread out during the dusk and dark hours, but form dense shoals or stay deeper at full daylight; probably as a predator avoidance tactic [36]. Considering that social and territorial behaviours were found to have genetic bases in related species of *Salvelinus* [37], we predicted that PL charr would display higher average values in social behaviours than SB charr. We also predicted reduced among-individual differences (low personality effect) in PL as selection would have depleted the genetic variation related to these traits. Moreover, because bold individuals appear to have lower propensities for social behaviours [38,39], we expected PL charr to have reduced average values as well as reduced among-individual differences in boldness as a response to selection for social personality types. In contrast, the high physical complexity of the habitats occupied by SB charr may not only relax the selection on social traits but also favour the establishment of a high diversity of personality types, for example by enabling bolder SB charr that are more likely to move across foraging areas or feeding away from shelters to thrive with shyer individuals that tend to stay within sheltered areas (e.g. fissures and restricted spaces between boulders).

**Table 1.** Predictions for the patterns of divergence between the two morphs regarding the three aspects of behavioral variation described in the text. PL: Planktivorous charr, SB: Small-benthic charr. The expected behavioural changes are not absolute, but relative to the two morphs.

| Morph | Habitat characteristics | Predictions: Sociality |  | Predictions: Boldness |  | 3. Trait correlation |
| --- | --- | --- | --- | --- | --- | --- |
|  |  | 1. Average trait value | 2. Personality effect | 1. Average trait value | 2. Personality effect |  |
| PL | - Open water, no shelter<br>- High spatio-temporal variations in prey availability | **High:** Response to selection favouring shoal formation | **Low:** Selection for social individuals reduces the diversity in personality types | **Low:** Selection against bolder individuals with low social propensity | **Low:** Reduced individual variation resulting from selection for social types | **Negative** (if enough among-individual variation) |
| SB | - Highly diverse physical structures<br>- Stable food resources | **Low:** Solitary/territorial life advantageous | **Low:** Sociality traits not relevant | **High:** Reduced selection against proactive behavioural types | **High:** Habitat heterogeneity maintaining individual polymorphism | **Null:** Shy individuals disadvantaged if social |

We raised individuals from pure-morph and hybrid crosses in common garden conditions to characterize the range of behavioural differences between the types of crosses regarding the three different aspects of variation. We expected the importance of the boldness and sociality traits in this case of divergence to be revealed by genetically based differences between SB and PL individuals according to the predictions described above. Moreover, because the merging of two diverging genomes often produces either maladaptive intermediate or transgressive hybrid traits[40,41], genetically-based behaviour variations in hybrids falling outside of the range of the two morphs could reveal whether they would be selected against.

## Methods

### Study system

Thingvallavatn is Iceland’s largest lake, with an area of 84km^2^ and a mean depth of 34 meters. The lake sits in a graben of the Mid-Atlantic ridge and was formed following the last glacial retreat about 10,000 years ago [42]. The physical structure of the lake is characterized by a wide pelagic zone and three major benthic habitats, being a “stony littoral” zone (0-10m deep) composed of a spatially complex lava substrate with loose stones, crevasses and interstitial spaces, a densely vegetated zone of *Nitella opaca* algae (10-20m deep), and a profundal zone (25 m and deeper) where the bottom is covered by a diatomic gyttja substrate [43]. The four morphs of Arctic charr (the planktivorous, the piscivorous, the large-benthic and the small-benthic), differ in habitat use, diet, head and body morphology, life-history and parasitism [30,43,44], and constitute at least three genetically differentiated populations (the status of piscivorous charr remains unresolved) [45]. All four morphs are completely sympatric, although coalescent models are consistent with scenarios involving short periods of geographic isolation between PL and SB charr [32]. The two morphs of our study overlap in their spawning seasons (SB: August-November, PL: September-November) [31] but show genetically-based differences in head shape [46], growth patterns [47] and foraging strategies [48]. Morphological differences between these two morphs are manifested early during development [46], but can be affected by limited plastic changes later in life [49]. The young of the year of the two morphs are believed to use the same habitat, the surf zone (0-1m deep), from the onset of active feeding in spring until the PL-charr shift towards deeper pelagic and epibenthic zones during summer [50].

### Field sampling of parental specimens and offspring rearing

We collected adult specimens in October 2017 by laying gillnets overnight on a spawning site used by the two morphs (Svínanesvík, 64°11’24.6”N; 21°05’40.5”W). We crossed the gametes of 18 ripe specimens as soon as they were brought ashore to generate nine full-sibling families of pure-morph (Female x Male parents : PLxPL and SBxSB) and hybrid crosses (PLxSB and SBxPL, see the crossing design in Table S1). The eggs were incubated in a single EWOS hatching tray (EWOS, Norway) at 4.1±0.2°C in the aquaculture facilities of Hólar University, Sauðárkrókur, Iceland. Hatching occurred in January 2018 and 20 to 40 free-swimming embryos per family were moved to single-individual cells in a common water flow-through tank on their hatching day (when 50% of their eggs clutch was hatched). Soon before the onset of active feeding (ca. 530 degree day, March 2018), we replaced the cells by 22cl identifiable, perforated and transparent cups allowing the exchange of olfactory cues as well as visual contact between individuals. At the same time, groups of ca. 20 fish that were not selected for the rearing experiment were moved into family-specific containers. These fish were used as test shoals for the experiment on sociality. All the fish were fed *ad libitum* with aquaculture pellets on a daily basis.

### Behavioural experiments

We conducted two types of experimental tests during a five-month period after hatching (May 2018, ca. 1100 degree days). In the wild this corresponds to the period when the juveniles of both morphs stay in the littoral zone, before PL charr shift towards deeper habitats [43]. The first test aimed at assessing the position of each fish along boldness/shyness axis (“boldness test”). The second test quantified the sociality of the same individuals (“sociality test”). Both tests relied on the video-tracking of a focal individual using the software Ethovision XT 8.5 (Noldus Information Technology, The Netherlands). 93 fish raised in the cups were used as focal individuals (37 PLxPL charr, 15 SBxSB charr, 23 and 16 F_1_ hybrids of PLXSB and SBxPL maternal origin, respectively).

The boldness test consisted of an open field test (OFT) with shelter [51,52]. This setup was composed of a 40×30×25 cm arena which was filled with ten liters of water from the tap of the raising tray (Figure 1b, Figure S1). The bottom left corner of each compartment contained an opaque white PVC shelter box (11×6.5×6cm) closed by a vertical sliding trapdoor. The area was divided into four virtual zones in relation to the shelter, the entrance zone of the shelter, a marginal zone and a central zone of the arena deemed to be the area of high risk. The width of the marginal zone was defined as twice the body-length of the focal fish. Our assumption on the spatial variation in risk level was based on thigmotaxis (the aversion for locations away from vertical surfaces), a concept commonly used in studies on boldness and anxiety [38]. The test started by introducing the focal fish into the shelter from the upper side through a two-centimeter-wide aperture, immediately sealed with a lid after the introduction. The trap door was gently opened after a five-minute acclimatization period and a 20-minute video-recording trial was simultaneously initiated. Twelve behavioral variables were extracted from the video output (Table S2). The test was repeated twice for every individual with a seven-day interval between trials in order to capture the behavioural variation related to both within- and among-individual differences. At the end of the first trial, a lateral view photograph of the left side of the specimen was taken for morphometric purpose using a down-facing fixed camera (Canon EOS 650D with a 100mm macro lens) before the fish was returned to the rearing tray.

The sociality tests were started one week later using an 80×30×15 cm arena divided in three compartments (Figure 2b, Figure S1). A central compartment (10×30 cm) contained the focal fish and was separated from two side compartments (10×30 cm) by transparent acrylic walls. These walls were perforated to allow transfer of chemical cues while preventing the fish from moving between compartments. A “start box” made of a vertical cylinder (10.5 cm high × 10.5cm inner-side diameter) was placed in the middle of the arena. Five fish of the same type of cross as the focal individual and raised in group since hatching were placed together in one of the two side compartments. The focal fish was introduced in the start box through a two-centimeter diameter door on the upper side. The start box was removed after an acclimatization period of five minutes and a 20-minute video record was initiated.

As with the experiment on boldness, two replicate trials were conducted for each fish, one week apart, but the side-compartment containing the group of congeners was alternated between the two rounds. The water of the arenas was always renewed between observations to mitigate the presence of chemical cues from the previous focal fish as well as temperature changes. In order to minimize stress, the fish were transported to the experimental room inside covered buckets filled with the same water as the common-garden setup. The trials were recorded using a video camera (IMAGING SOURCE DMK 21AU04, 640×480 pixels) placed 180 centimeters above the center of the multi-arena setups and operated with the software IC-capture 2.4 (with a frame rate of 30 Hz). After the last trial, each fish was euthanised with an overdosage of 2-Phenoxyethanol [53] and weighted to obtain the wet body mass.

### Statistical analyses

We first developed a straightforward boldness/shyness index based on the twelve variables recorded during the open-field tests by conducting an Exploratory Factor Analysis (Table S2). Briefly, this method characterises unobserved “latent” variables associated to sets of correlated observed variables [54]. We ran the analysis using the Maximum Likelihood method available in the R package psych [55] and identified two latent variables. One latent variable was related to observed variables describing a classical “boldness/shyness” axis as well as exploratory tendencies (e.g. travelled distance, velocity), and will hereafter be referred to as the *boldness* trait (Figure S2, Figure 2b). The second latent variable regrouped the observed variables related to the use of the entrance of the shelter (e.g. entries and exits frequencies, time spent in the entrance zone). Because this variable mostly describes the actions of the fish near the aperture of the shelter, we name it the *doormat* trait. For validation, we also reduced our dataset using Principal Component Analyses (PCA), a classical alternative method to derive personality scores [56,57]. The resulting first two principal components reflected the boldness and doormat traits (Figures S4).

In order to account for the confounding effect the physical condition of the fish may have on their behavioural response, we extracted individual indexes of body condition as residuals from a regression of the wet weight of the specimen over its standard length [58] (Figure S4, Table S3). This value was extracted from the morphometric photographs using the R packages StereoMorph and Geomoporph [59,60].

We used Multi-response Linear Mixed Models [61] to test whether the types of cross differed in the three behavioural aspects (1: average trait value, 2: individual consistency, and 3: trait covariance), by using as a multiple response the values of the three traits (boldness, doormat, *sociality*), mean centred and scaled by their respective standard deviations. We studied behavioural aspect 1 by assessing the importance of the type of cross as a fixed effect in a multi-response model containing standard length, body condition and trial number as covariates. The identity of the specimen and its family were added as random variables. We assessed the importance of the effect of the type cross relative to the total amount of behavioural variation by adapting the marginalized determination coefficient (*R*^2^_*m*_) from [62] :

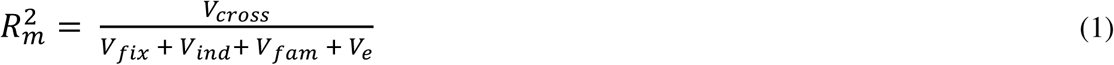

where *V*_*cross*_ and *V*_*fix*_ are the variances calculated from the fixed effect component referring to the type of cross alone and to all fixed effects, respectively [63]. *V*_*ind*_ and *V*_*fam*_ are the variance components associated with the differences between the intercept of individuals and of families, respectively, and *V*_*e*_ is the residual variance (within-individual variance).

The differences between types of crosses related to the second and the third behavioural aspects (consistent individual differences and behaviour syndrome) were assessed by extracting the variance and covariance components of three separate models (one per type of cross). These models contained the standard length, body condition, trial number and family identity as fixed effects while the individual identity was set as a random variable. From these three models, we assessed the amount of consistent differences between individuals in each trait and in each type of cross by calculating their adjusted repeatability (*R*). The adjusted repeatability controls for confounding factors (here body condition, size, trial number and family) and was calculated using the formulation from [64]:

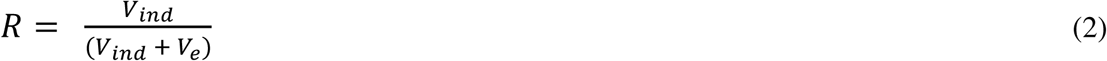

Finally, we tested for differences in correlations between traits (an important component of behavioural syndromes, aspect 3) among types of cross by comparing the between-trait correlation coefficients extracted from the variance-covariance matrices of the three models. In order to gain statistical power, the two categories of reciprocal hybrids were pooled for all models.

We fitted all the models under a Bayesian framework using Markov Chain Monte Carlo (MCMC) methods as implemented in the package MCMCglmm [61]. We specified weakly informative priors (V_0family_ = 1, V_0ind_ and V_0res_ = identity matrix *I*_3_, *nu* = 0.002) and determined the number of iterations allowing model convergence through the examination of trace plot, posterior density plots and effective sample sizes. Inferences were made by comparing the posterior mode estimates and 95% Highest Posterior Density Credible intervals (95% CrI) between the type of crosses (and in relation to the zero baseline for the significance of *R* estimates).

## Results

We found that the boldness scores tended to be lower and the average distances to conspecifics tended to be higher (lower sociality) in SBxSB offspring and hybrids than in PLxPL offspring (Figure 3, Table S4). The effect of the type of cross, however explained a negligible proportion of the total variation (R^2^_m_ : posterior mode [95% CrI] = 0.01 [0.00-0.03]). These results were also observed as limited trends in the graphical representations of the reaction-norms of each trait (Figure S5) but were also non-significant when employing separate linear mixed models with a single, non-scaled trait as a response (Table S5).

**Figure 3.**
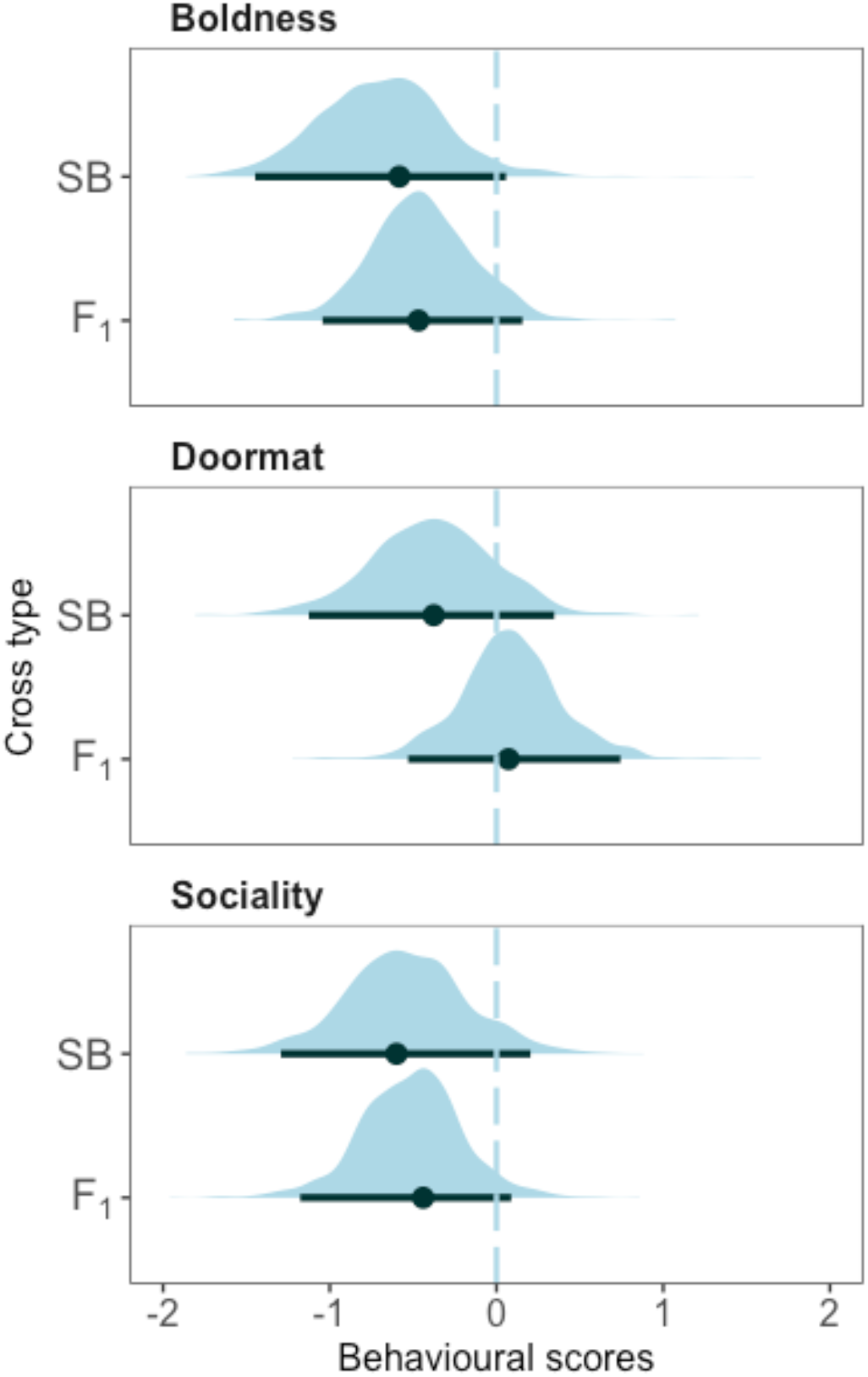
Posterior distribution, posterior mode and 95% HPD intervals of the fixed effect “type of cross” (SBxSB offspring: SB, hybrids: F_1_) on each trait from the multiple-response model. The PL-charr constitutes the baseline (dashed-line). The scores are mean-centred and scaled by unit of standard deviation. Behavioural scores for Boldness and Doormats: scores of latent variables for a Factor Analysis; for Sociality: inverse of the average distance to conspecific (cm).

The posterior modes of the repeatability estimates were high (> 0.5) in SBxSB and PLxPL offspring for the boldness and the doormat traits, and were supported by lower limits of 95% CrI with high values (Figure 4a, Table S6). These results indicate high levels of individual consistency (high “personality effect”) in these two traits for both morphs. This effect was not observed for the sociality trait, the posterior modes of repeatability estimates being low in the two morphs. In contrast, the posterior modes of repeatability estimates were medium to low for all traits in hybrids; the posterior density of these estimates furthermore included or were close to zero. The repeatability estimates of all traits did not appear to differ among the pure-bred offspring. Considering the posterior density of these estimates, we can say with good confidence that the hybrids showed lower individual consistency in boldness than at least the SBxSB-offspring. The repeatability estimates of doormat were also lower in the hybrids than in the PLxPL offspring.

**Figure 4.**
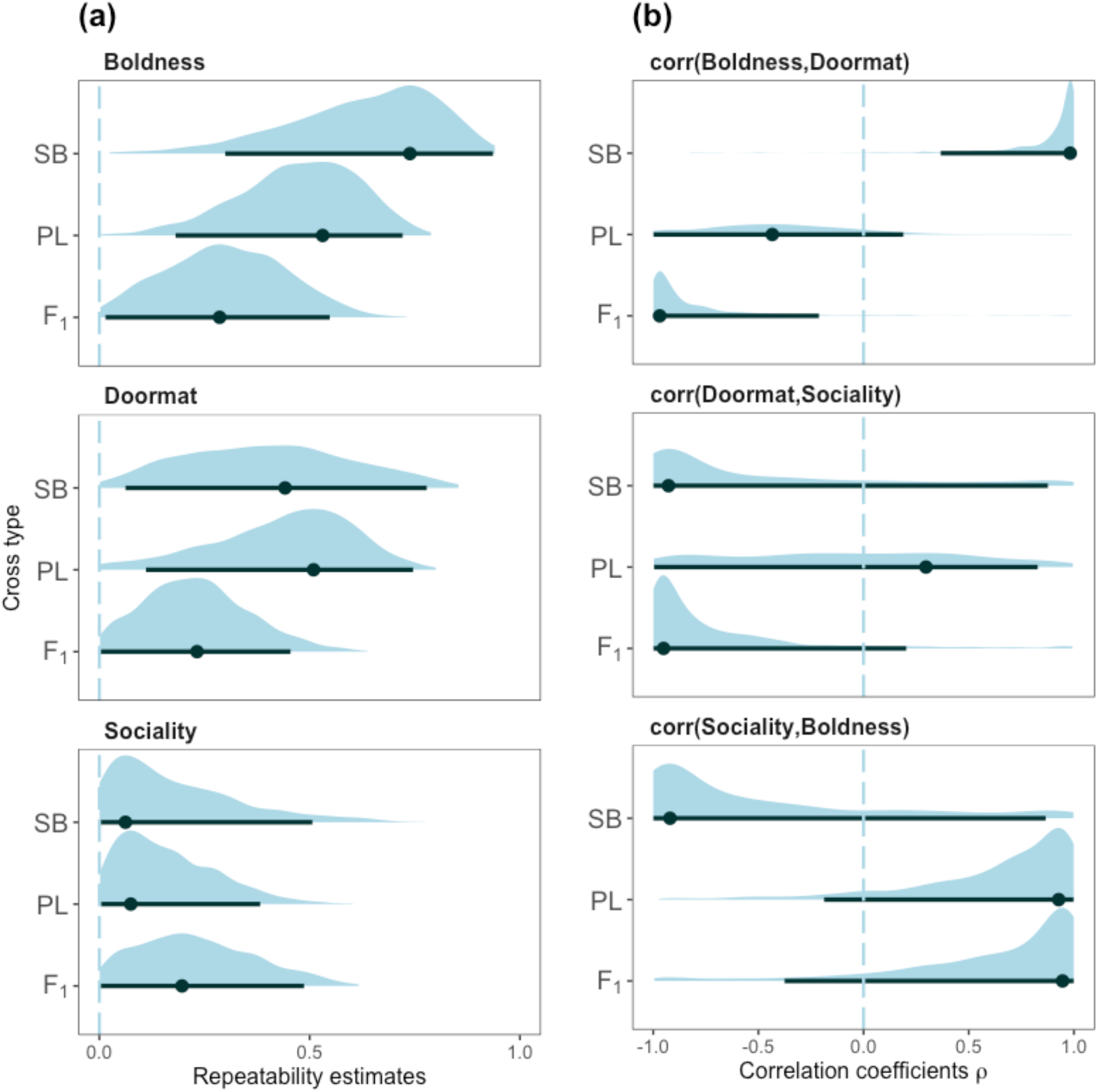
Posterior distributions, posterior mode and 95% Credibility Intervals of (a) the repeatability estimates for the tested trait and (b) of the Pearson correlation coefficients between traits at the individual level (behaviour syndrome) for each type of cross. Types of cross: SBxSB offspring (SB); PLxPL offspring (PL); hybrids (F_1_).

We also observed differences in the posterior estimates of trait covariances (i.e. behavioural syndromes) between the types of crosses (Figure 4b, Table S6). Among SBxSB offspring, the posterior modes of the correlation estimates showed a positive correlation between boldness and doormat behaviour whereas the corresponding correlation among PLxPL offspring was negative. The correlation estimates between doormat and sociality were mostly negative in SBxSB offspring while no trend was observed in PLxPL offspring. The correlation estimates between sociality and boldness also differed between pure-morph offspring. The corresponding estimates were strongly positive in the PLxPL offspring and higher than in the SBxSB offspring where the estimates were mostly negative.

Differences were also observed in the trait correlations in hybrids compared to those of the pure-bred offspring (Figure 4b, Table S6). Hybrids showed a negative correlation between boldness and doormat in hybrids that differed from the negative estimate of SBxSB offspring, and which tended to be even more negative than those of the PLxPL offspring. The covariance estimates between boldness and sociality were also similar between the hybrids and the SBxsB offspring. However, the correlation between sociality and boldness was positive in hybrid and did not differ from the one of the PLxPL charr. These estimates also differed from the negative correlation between sociality and boldness in SBxSB charr.

## Discussion

Like any trait with genetic bases, behavioural traits undergoing adaptive divergence can be revealed through common garden experiments [65]. Our hypothesis that adaptive divergence can act on the three aspects of behavioural variation studied here is partially supported by the complexity of the results from such experiments. First, our data provides little support to our predictions that the two morphs differ in average values of their behaviour traits. Second, the two morphs showed the same pattern in repeatability of the three traits. They displayed high levels of repeatability in boldness and doormat, while no repeatability was observed in social behaviour. Third, contrasting behavioural syndromes were found between the two morphs as seen in the opposite patterns of covariance between boldness and doormat, as well as between boldness and sociality.

The hybrids differed from the two morphs in a complex way. The hybrids showed reduced repeatability in boldness and doormat behaviour compared to pure-morph offspring. They showed similar patterns as PL offspring in the covariations of boldness-doormat and boldness-sociality, the two set of covarying traits that significantly differed between the two pure-morph offspring. Conversely, the hybrids tended to be more similar to SB-offspring in the behavioural syndrome involving doormat and sociality.

### 1. Personality as a driver of adaptive divergence

Contrary to our predictions, the two morphs did not differ in average values (our first aspect of behavioural variation) of boldness- and sociality, but both had high repeatability estimates (second aspect of variation) for those two traits. This high consistency may reflect key characteristics of the respective foraging environment of each morph and can be interpreted in light of individual specialization [66]. Differences in the degree of individual specialization can emerge between habitats. For example, in lake populations of Eurasian perch (*Perca fluviatilis*) individuals utilizing the littoral zone have a more specialized diet than their pelagic congeners [67]. Animal personality can be linked to individual specialisation as a consequence of individuals developing alternative strategies (*e.g.* for energy acquisition) [3,5]. In our system, these high levels of consistent differences between individual PL juveniles probably reflects more complex evolutionary responses than expected (Table 1) but may be related to conditions offering an advantage for specialized behavioural types in the pelagic habitat as well. Different personality types in the PL-offspring could have emerged as a result of more complex ecological characteristics than expected in the pelagic habitats, and where different strategies related to resource acquisition and/or “predation risk *versus* energy gain” trade-offs [68] yield similar fitness outcomes. At the height of the productive season PL charr mainly prey on zooplankton, especially *Daphnia longispina* and *Cyclops abyssorum* [69], two species with high spatiotemporal variations in availability. Partly this results from diel migration cycles of the zooplankton, but variations in horizontal distribution may also be important [70]. Such spatiotemporal complexity of food resources, probably coupled with the predation risk inherent to open habitats, may generate energy acquisition trade-offs that affect foraging strategies of PL charr. This is supported by the experimental observations where naive PL offspring, when offered live Daphnia as food, appeared not to start feeding bellow a given threshold of prey density. Such restraint was not seen in SB-offspring [48].

The environmental conditions favouring the diversity of personality types can also be related to the social environment experienced by PL charr. For example, bolder zebrafish (*Danio rerio*) are more likely to be the dominant individuals [38] and such personality-related hierarchical structures can be expected in Arctic charr, as their food intake, dominance and swimming activity are mediated by common hormonal mechanisms [71]. Finally, the complexity of the social environment can also be intertwined with fitness trade-offs, as seen for example in sticklebacks, where bolder fish were more dominant, grew faster and occupied frontal positions of shoals where the foraging success is higher, but where the predation risk is also higher [39]. Similar patterns may be expected in the PL charr that both form dense shoals and are confronted to spatial-temporal heterogeneity in their food resources [70].

The observation of substantial but different personality syndromes (our third behavioural aspect) in both morphs was contrary to our predictions stating that SB charr should not show display covariance in the studied traits. Because the evolution of a behavioural syndrome can be related to situation-specific conflicts in the expressions of behavioural responses [4], these observations may also result from different energy acquisition trade-offs among benthic and limnetic habitats and/or imposed by the different social environments discussed above. These different covariance patterns can also be explained more generally as a response to two different regimes of correlational selection acting in each habitat [25]. Although the precise ecological and evolutionary factors responsible for such differences in trait covariance remain unclear, these findings suggest that SB- and PL-charr have undergone substantial adaptive divergence at the level of behavioural syndromes.

Because the fitness consequences of the behavioural variations we observed were not directly measured, we call for a cautious interpretation of our results. Caution is also warranted as the connections between personality and individual performance are nontrivial [72–74]. We however are confident that the behavioural types we observed can be interpreted as phenotypic values maximizing fitness in the different habitats. Based on more than thirty years of studies of the Arctic charr of Thingvallavatn it can be stated with good confidence that many of the morphological and life-history-related differences between PL and SB charr have evolved from diversifying selection related to trophic and non-trophic ecological factors [43,45,75]. One would therefore find it likely that in this particular case of divergence, behavioural traits would be affected in a similar way.

### 2. Hybrids behaviour and implications in speciation

The merging of diverging genomes often results in transgressive or intermediate values of polygenic traits [40], and transgressive or intermediate behaviours in hybrids have recently been proposed as an overlooked source of post-zygotic reproductive isolation between diverging populations [18,76]. Although the hybrids from our experiment did not differ the two morphs in their average behavioural responses, they tend to show reduced repeatability in the traits for which the two morphs show high level of consistent individual differences. Repeatability may therefore be affected in the same way as non-behavioural characters by hybrid breakdown (*i.e.* deficiencies resulting from the negative genetic interactions of the incompatible alleles from diverging genomes [77]).

Furthermore, hybrids of the two morphs show a particular pattern of trait covariance, as in traits where SB and PL charr differ in their behaviour syndromes, the hybrid phenotype in one case follows the SB-charr but in another the PL-charr. While theoretical views suggest that breakdowns through hybridization in the genomic architecture responsible for trait correlations may generate transgressive characters [78], the hybrid phenotypes observed here show a complex picture in which the different trait covariances may be affected by different mechanisms. The absence of intermediate hybrid phenotypes suggests that different, non-additive genetic effects are responsible for the development of such syndromes. Such results are consistent with the observation that the development of foraging-related personality traits in nine-spine stickleback (*Pungitius pungitius*) is dependent of non-additive genetic factors [65], although trait covariances were not investigated in this study.

Whatever the proximate mechanisms, the complex covariance patterns observed in the hybrid charr result in an idiosyncratic phenotype that may not perform as well as pure-morph individuals in their respective ecological niches. Although novel phenotypes may facilitate the colonisation of still-unoccupied niches [40], such advantage appear to be unlikely in the contemporary state of our system. Hybrids between PL and SB charr are indeed virtually non-existent in the wild, at least at the adult stage, in spite of what appear to be ample opportunities for the two morphs to interbreed [31].

Together with reduced individual consistency, the singular patterns of personality syndromes may be a source of selection against hybrids. While the extent to which such selection constitutes a reproductive barrier (especially in relation to physical traits) and how much it contributes to the total level of reproductive isolation between the two morphs remains to be ascertained. The present findings suggest that studying simultaneously several aspects of behavioural variation can uncover nonintuitive patterns of adaptive divergence.

## Acknowledgement

We are very thankful to Alia Desclos for her contribution in operating the experimental test and processing the videos and her help for the maintenance of the raising setup. We thank Skúli Skúlason for his comments of the manuscripts, Zophonías O. Jónsson, the members of the “Arctic charr and salmonid” lab of the university of Iceland and the farmer Jóhann Jónsson for their help during the sampling, and Camille Leblanc, Bjarni K. Kristjánsson, and Neil Metcalfe for constructive discussions on the experimental design and the conceptual aspects of the study. We thank Kári H. Árnason, Rakel Þorbjörnsdóttir and Christian Beuvard for the organisation and the maintenance of the growing facility.

## Founding

This work was entirely funded by the Icelandic Centre of Research, RANNÍS (Icelandic Research Fund grant no.173802-051).

## Ethical statement

Fishing was conducted with the permissions of the owner of the farm of Mjóanes and of the Thingvellir National Park commission. Ethics committee approvals for research project are not required by the Icelandic regulation (Act No. 55/2013 on Animal Welfare). However, the experimental work was conducted at the Hólar University Aquaculture Research Station, an institute owning an operational license in line with the Icelandic law on Aquaculture (Law No. 71/2018), which includes closes of best practices for animal care and experimental work.

## Authors’ contribution

QJBH conceived the study, operated the common-garden setup, conducted the data collection, carried-out the statistical analyses, and drafted the manuscript. DB designed the open-field tests, organised the data collection, supervised the video processing steps and contributed to the writing. SSS coordinated the field work, produced these embryos and critically revised the manuscript. MBM provided guidance during the data analyses, contributed to the biological interpretations of the results and reviewed the manuscript. KHK generated the crossing design, produced the embryos, planned their transfer and maintenance to the aquaculture facility and critically revised the manuscript. All authors gave final approval for publication and agree to be accountable for the work therein.

