## Supplemental figures and tables for "Animal personality adds complexity to the processes of adaptive divergence and speciation"

### Supplementary Material – Additional results.

**Table S1.** Crossing design of the rearing experiment and number of individuals used for the personality tests. Type of cross: Female gamete x Male gamete. Parents were used for crosses only once (no split families).

| Type of cross | Number of families | Number of individuals |
|---|---|---|
| **PLxPL** | 2 | 37 |
| **SBxSB** | 2 | 15 |
| **PLxSB** | 3 | 23 |
| **SBxPL** | 2 | 18 |

**Table S2.** Variables measured during the open-field test with shelter.

| *Variable name* | *Description* |
|---|---|
| Angular speed | Absolute angular velocity of the fish (degrees per second), calculated using the following equation: $V_{angn} = RTA_n(t_n - t_{n-1})^{-1}$ where $RTA_n$ is the relative turn angle of the sampled position *n*, and *t* the sampling time. The rate of change in direction was unsigned. The turn angle was calculated as the difference between two subsequent values for the direction of the head. This variable quantify the extent to which the fish was turning around itself and is an index of swimming path complexity. High $V_{angn}$ values can be referred to high levels of vigilance (Benhaïm et al., 2012). |
| Arena center - Freq | Number of events involving the fish entering the central zone |
| Arena center - Time | Total time spent in the central zone (seconds) |
| Arena edges - Time | Time spent within the marginal zone of the arena |
| Average distance to shelter | Mean distance between the fish and the shelter (cm) |
| Latency to leave shelter | Time lagg between the removal of the trapdoor and the first exit of the fish (seconds) |
| Shelter - Freq | Number of events involving the fish returning to the shelter |
| Shelter - Time | Total time spent inside the shelter (seconds) |
| Shelter entrance - Frequency | Number of events involving the fish entering the shelter |
| Shelter entrance - Time | Total time spend in the entrance zone infrom of the shelter |
| Total distance | Total distance swam (cm) |
| Velocity | Mean velotcity (body length swam per second) |

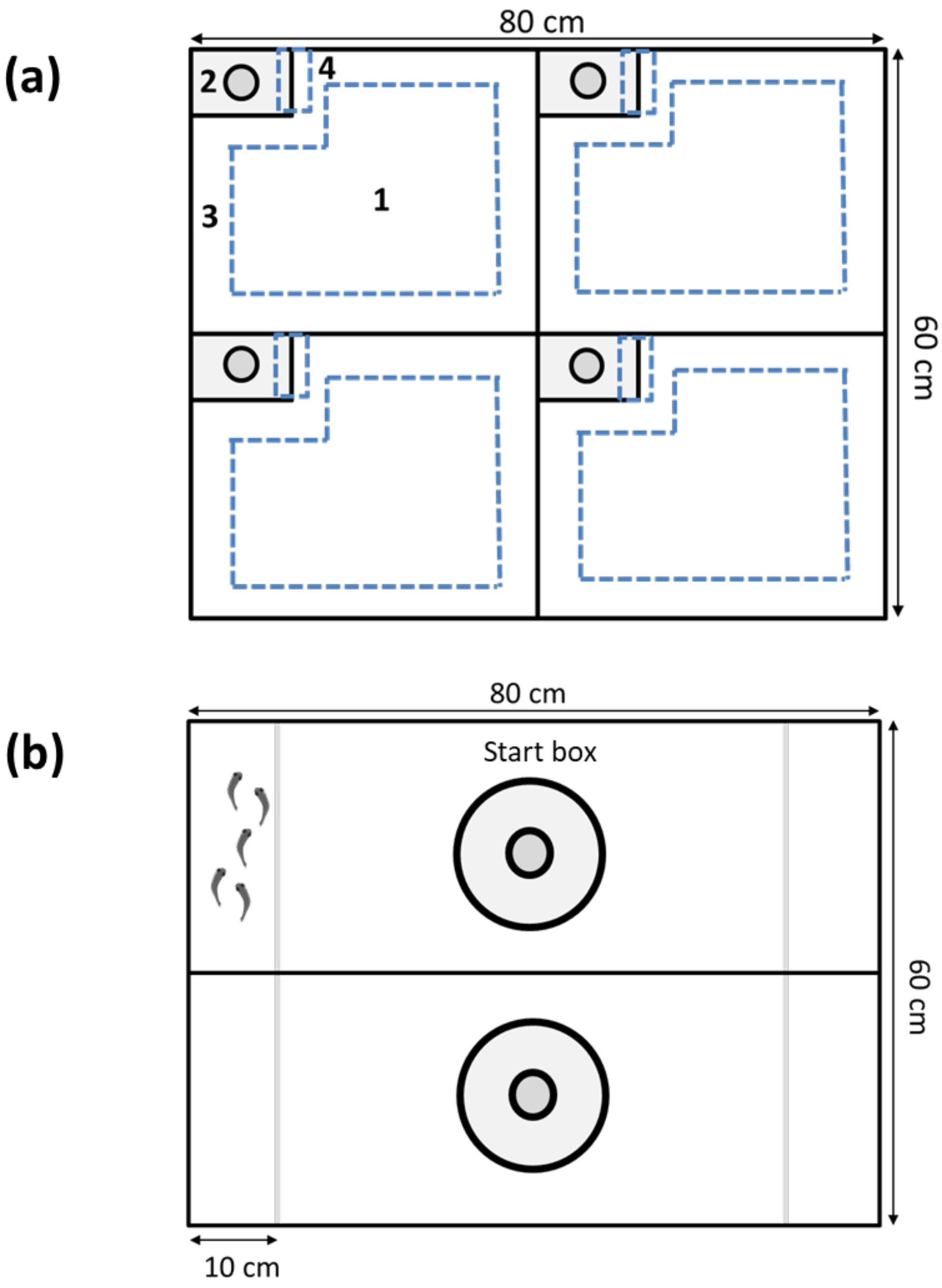


**Figure S1.** Dimensions of the multi-arena setups used for the boldness (a) and the sociality (b) tests (aerial views). The different virtual zones, delimited by the dashed lines are: 1:the center zone, 2: the shelter, 3: the marginal zone, 4: the entrance area. Transparent walls are represented by grey continuous lines. A group of congeners is depicted in the upper-left compartment in (b). The darker circles are the lit of the shelters and the start boxes, through which the focal fish was introduced.

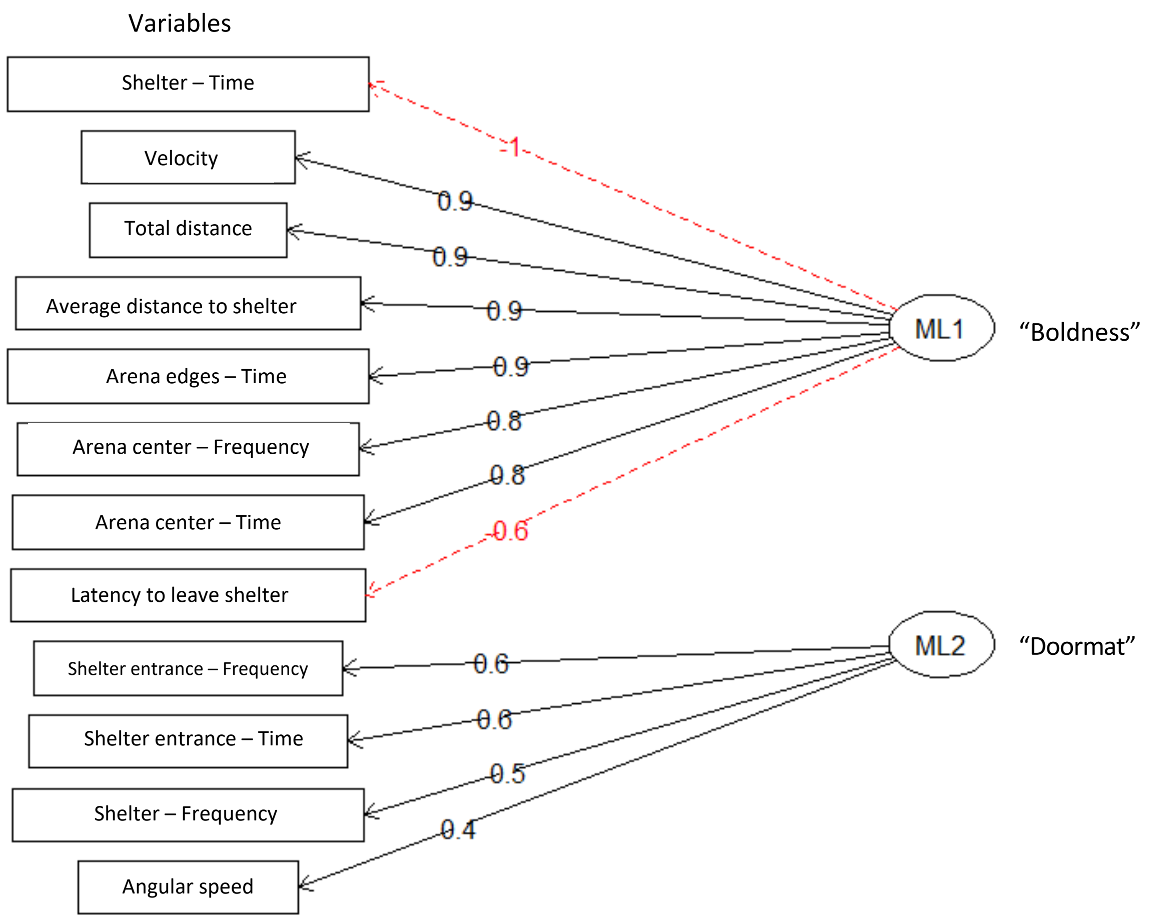


**Figure S2.** The two latent variables (ML1 and ML2) generated through factor analyses of the variables from the *boldness* test. Figures on arrows: Factor loadings. Continuous lines: positive correlation; dashed lines: negative correlations. ML1 and ML2 are referred in the subsequent analyses as the "boldness" and the "doormat" trait. respectively. Refer to Table S1 for the description of the variables.

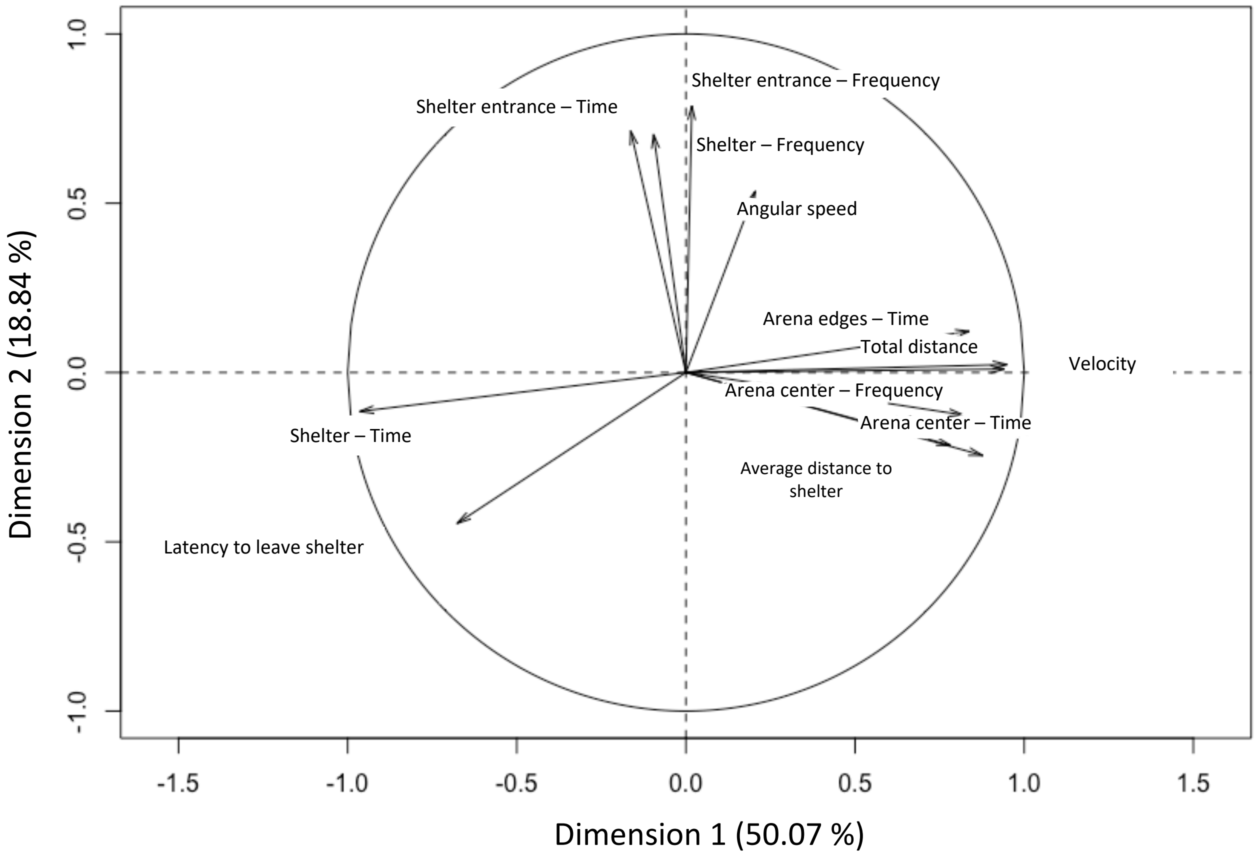


**Figure S3.** Principal components map of the 12 variables for the boldness test. Dimension 1 is comparable to the *boldness* latent variable from the Factor Analysis while Dimension 2 is similar to the *doormat* variable.

*1. Body condition.*

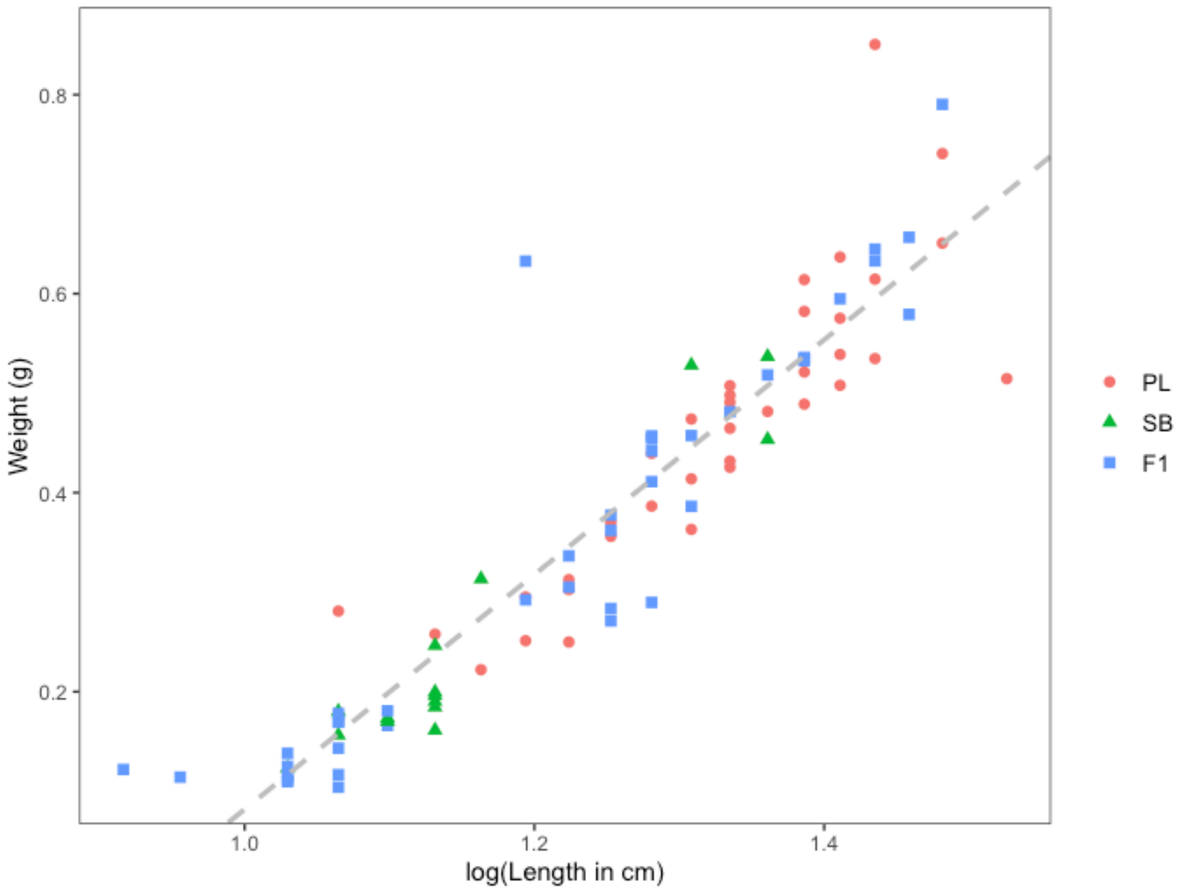


**Figure S4.** Weight of each specimen regressed over size. The residuals were used in subsequent model as estimates of body condition. PL: Planctivorous. SB: Small-benthic char. F1: $F_1$ hybrids. Dashed line: Regression line according of Model number 1. Table S1.

**Table S3.** Table of variance of a reduced (1) and full (2) model of variation of weight with size. The residuals of the reduced model were used as estimated of body condition. Adjusted $R^2$ = 0.86 ; 0.86 ; df = 91 . 87 ; F = 569.5 ; 111.1 for Model number 1 and 2. respectively.

| Model | Formula | Coefficient | Estimate | Std. Error | t-value | *p*-value |
|---|---|---|---|---|---|---|
| **1** | Weight ~ 1 + log(length) | Intercept | -1.10 | 0.06 | -17.62 | <0.01 |
| | | log(length) | 1.18 | 0.05 | 23.86 | <0.01 |
| **2** | Weight ~ log(length) + cross + log(length) x cross | Intercept (crossPL) | -1.12 | 0.15 | -7.54 | <0.01 |
| | | log(length) | 1.19 | 0.11 | 10.69 | <0.01 |
| | | crossSB | -0.10 | 0.26 | -0.38 | 0.71 |
| | | crossF1 | 0.05 | 0.17 | 0.28 | 0.78 |
| | | log(length) x crossSB | 0.08 | 0.21 | 0.37 | 0.71 |
| | | log(length) x crossF$_1$ | -0.03 | 0.13 | -0.21 | 0.84 |

*2. Graphical explorations with Reaction norms.*

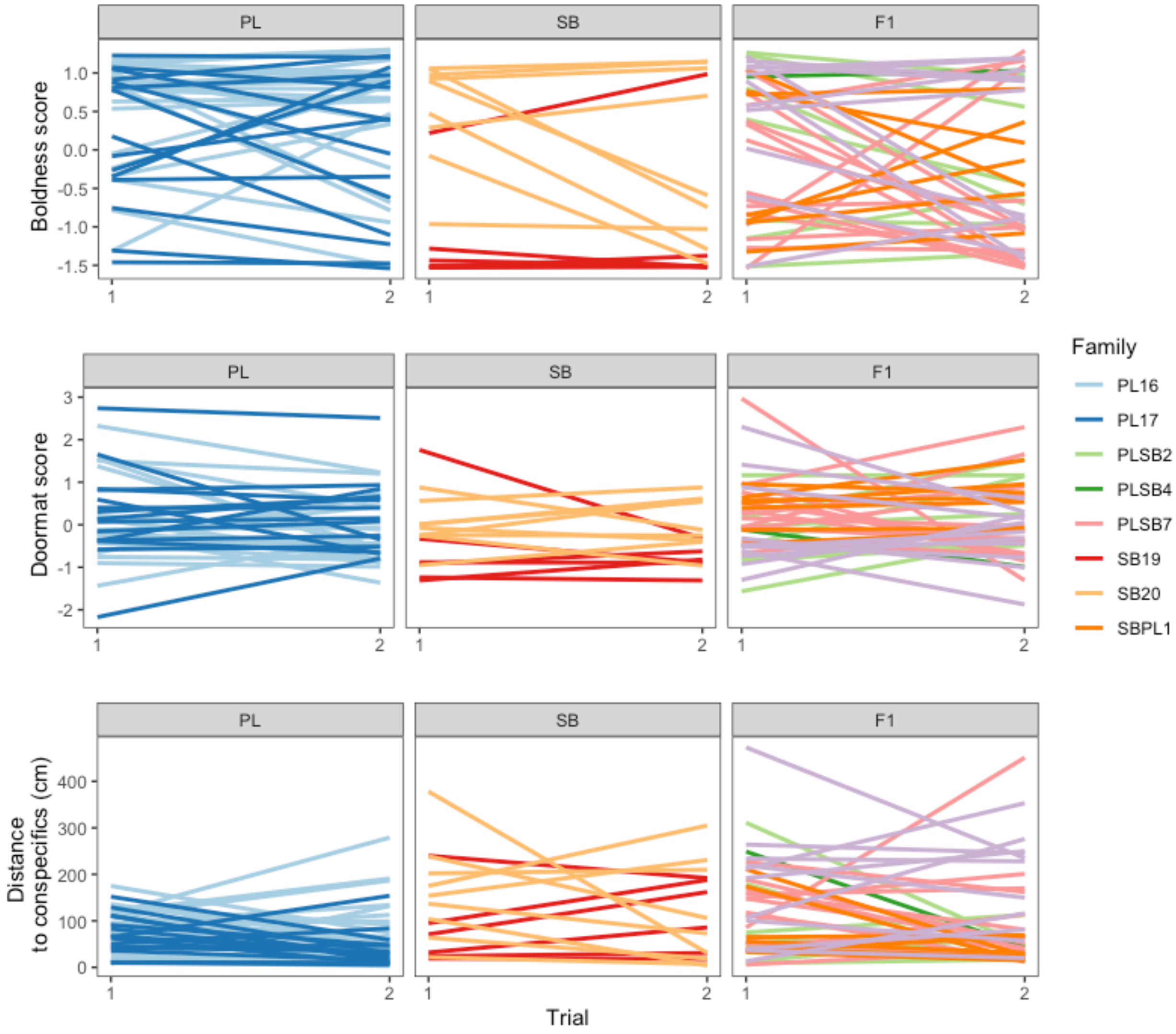


**Figure S5.** Reaction norms of the behavioural response of each type cross Each line refers links the response of one individual between two repetitions of the behavioural test involved. PL: pure-morph planktivorous cross. SB: pure-morph small-benthic cross. The average distance to conspecifics (Lower panel) constitutes the negative of the Sociality trait.

#### 3. *Multiple-response models.*

**Table S4.** Summary table of Model 1. the multiple-response mixed-effect model considering "boldness". "doormat" and "sociality" as a multiple-response. $V_{ind}$ : Inter-individual variance component. Non-overlapping 95% High Posterior Density Credible Interval (95% CrI) are use to detect significant differences between effects. Refer to Figure 3b in the main text for clearer interpretations. Cross type-PL constitutes de baseline. Sociality is here interpreted as the average distance to conspecifics (higher values related to less social individuals).

| Effect | Posterior mode | 95% CrI |
|---|---|---|
| *Fixed effects* | | |
| Boldness | 0.27 | -0.23 - 0.77 |
| Doormat | -0.03 | -0.47 - 0.48 |
| Sociality | -0.38 | -0.91 - 0.10 |
| Length | 0.00 | -0.11 - 0.14 |
| Boldness x Trial | -0.12 | -0.22 - 0.00 |
| Doormat x Trial | -0.11 | -0.18 - 0.06 |
| Sociality x Trial | -0.08 | -0.23 - 0.02 |
| Boldness x Cross type-SB | -0.56 | -1.52 - 0.03 |
| Doormat x Cross type-SB | -0.39 | -1.14 - 0.35 |
| Sociality x Cross type-SB | 0.34 | -0.22 - 1.29 |
| Boldness x Cross type-$F_1$ | -0.61 | -1.04 - 0.17 |
| Doormat x Cross type-$F_1$ | 0.08 | -0.53 - 0.74 |
| Sociality x Cross type-$F_1$ | 0.45 | -0.09 - 1.18 |
| Boldness x Body condition | 0.49 | -2.66 - 2.51 |
| Doormat x Body condition | -0.67 | -3.56 - 1.62 |
| Sociality x Body condition | 1.25 | -1.12 - 3.34 |
| *Random Effects* | | |
| Family | 0.03 | 0.00 - 0.26 |
| Boldness ($V_{ind}$) | 0.33 | 0.15 - 0.63 |
| Doormat x Boldness ($V_{ind}$) | -0.19 | -0.36 - 0.00 |
| Sociality x Boldness ($V_{ind}$) | 0.04 | -0.10 - 0.23 |
| Doormat ($V_{ind}$) | 0.30 | 0.05 - 0.56 |
| Sociality x Doormat ($V_{ind}$) | -0.16 | -0.33 - 0.03 |
| Sociality ($V_{ind}$) | 0.21 | 0.00 - 0.40 |
| *Residuals* | | - |
| Boldness | 0.46 | 0.37 - 0.69 |
| Doormat x Boldness | 0.06 | -0.06 - 0.22 |
| Sociality x Boldness | 0.06 | -0.08 - 0.22 |
| Doormat | 0.69 | 0.51 - 0.97 |
| Sociality x Doormat | -0.05 | -0.28 - 0.08 |
| Sociality | 0.69 | 0.51 - 0.95 |

**Table S5.** Fixed effect structure of three seperate linear mixed-effect models with each trait as a response. Non-overlapping 95% High Posterior Density Credible Interval (95% CrI) are use to detect significant differences between effects. Sociality is here interpreted as the average distance to conspecifics (higher values related to less social individuals).

| Response | Effect | Posterior Mode | 95% CrI |
|---|---|---|---|
| **Boldness** | | | |
| | Cross Type-PL (Intercept) | 0.30 | -0.37 - 1.41 |
| | Body condition | 0.33 | -2.55 - 2.43 |
| | Trial | -0.06 | -0.43 - 0.21 |
| | Cross Type-SB | -0.20 | -1.85 - 1.03 |
| | Cross Type-Hybrid | 0.06 | -1.27 - 1.03 |
| | Trial x Cross Type-SB | -0.11 | -0.84 - 0.34 |
| | Trial x Cross Type-Hybrid | -0.24 | -0.69 - 0.20 |
| **Doormat** | | | |
| | Cross Type-PL (Intercept) | 0.34 | -0.22 - 1.08 |
| | Body condition | -0.88 | -3.02 - 1.37 |
| | Trial | -0.21 | -0.58 - 0.09 |
| | Cross Type-SB | -0.53 | -1.67 - 0.47 |
| | Cross Type-Hybrid | -0.23 | -1.00 - 0.71 |
| | Trial x Cross Type-SB | 0.08 | -0.36 - 0.78 |
| | Trial x Cross Type-Hybrid | 0.23 | -0.28 - 0.60 |
| **Sociality** | | | |
| | Cross Type-PL (Intercept) | 4.57 | 3.67 - 5.30 |
| | Body condition | 2.41 | -0.56 - 4.39 |
| | Trial | -0.38 | -0.77 - 0.00 |
| | Cross Type-SB | 0.26 | -0.96 - 1.80 |
| | Cross Type-Hybrid | 0.04 | -0.89 - 1.37 |
| | Trial x Cross Type-SB | 0.10 | -0.75 - 0.71 |
| | Trial x Cross Type-Hybrid | 0.08 | -0.43 - 0.68 |

**Table S6.** Posterior modes (Post. mode) and 95% High Posterior Density Credible Interval (95% CrI) of the variance-covariance components and repeatability estimates for the three multiple-response mixed models (one per morph). Refer to Figure 4 in the main text for clearer interpretations. COV(,) : covariance.

| | **Cross type** | | | | | | | | | | | |
|---|---|---|---|---|---|---|---|---|---|---|---|---|
| | Pure-PL | | | | Pure-SB | | | | $F_1$ | | | |
| | Post. mode | 95% CrI | | | Post. mode | 95% CrI | | | Post. mode | 95% CrI | | |
| **Among-individual variance-covariance** | | | | | | | | | | | | |
| Boldness | 0,43 | 0,14 | - | 0,98 | 0,38 | 0,06 | - | 1,93 | 0,30 | 0,00 | - | 0,66 |
| COV(Boldness,Doormat) | -0,23 | -0,55 | - | 0,14 | 0,28 | -0,08 | - | 1,51 | -0,23 | -0,50 | - | 0,01 |
| COV(Boldness,Sociality) | 0,18 | -0,10 | - | 0,45 | -0,08 | -0,75 | - | 0,43 | 0,14 | -0,11 | - | 0,43 |
| Doormat | 0,55 | 0,00 | - | 0,98 | 0,28 | 0,00 | - | 1,49 | 0,24 | 0,00 | - | 0,66 |
| COV(Doormat,Sociality) | -0,04 | -0,30 | - | 0,28 | -0,07 | -0,75 | - | 0,39 | -0,16 | -0,44 | - | 0,07 |
| Sociality | 0,08 | 0,00 | - | 0,45 | 0,01 | 0,00 | - | 0,79 | 0,18 | 0,00 | - | 0,58 |
| **Within-individual variance-covariance** | | | | | | | | | | | | |
| Boldness | 0,47 | 0,29 | - | 0,85 | 0,32 | 0,14 | - | 0,69 | 0,66 | 0,45 | - | 1,08 |
| COV(Boldness, Doormat) | 0,01 | -0,28 | - | 0,21 | -0,08 | -0,40 | - | 0,17 | 0,10 | -0,14 | - | 0,38 |
| COV(Boldness, Sociality) | -0,14 | -0,41 | - | 0,06 | 0,25 | -0,03 | - | 0,66 | 0,03 | -0,18 | - | 0,33 |
| Doormat | 0,59 | 0,34 | - | 1,02 | 0,52 | 0,31 | - | 1,26 | 0,82 | 0,58 | - | 1,29 |
| COV(Sociality, Doormat) | -0,11 | -0,33 | - | 0,23 | -0,14 | -0,77 | - | 0,11 | -0,14 | -0,42 | - | 0,10 |
| Sociality | 0,87 | 0,59 | - | 1,25 | 0,77 | 0,49 | - | 1,72 | 0,82 | 0,45 | - | 1,12 |
| **Repeatability** | | | | | | | | | | | | |
| Boldness | 0,44 | 0,18 | - | 0,72 | 0,72 | 0,30 | - | 0,94 | 0,24 | 0,01 | - | 0,55 |
| Doormat | 0,45 | 0,11 | - | 0,75 | 0,35 | 0,06 | - | 0,78 | 0,26 | 0,00 | - | 0,45 |
| Sociality | 0,07 | 0,00 | - | 0,38 | 0,01 | 0,00 | - | 0,51 | 0,27 | 0,00 | - | 0,49 |
